# The landscape of cellular clearance systems across human tissues and cell types is shaped by tissue-specific proteome needs

**DOI:** 10.1101/2024.08.26.609695

**Authors:** Ekaterina Vinogradov, Lior Ravkaie, Bar Edri, Juman Jubran, Anat Ben-Zvi, Esti Yeger-Lotem

## Abstract

Protein clearance is fundamental to proteome health. In eukaryotes, it is carried by two highly conserved proteolytic systems, the ubiquitin-proteasome system (UPS) and the autophagy-lysosome pathway (ALP). Despite their pivotal role, the basal organization of the human protein clearance systems across tissues and cell types remains uncharacterized. Here, we interrogated this organization using diverse omics datasets. Relative to other protein-coding genes, UPS and ALP genes were more widely expressed, encoded more housekeeping proteins, and were more essential for growth, in accordance with their fundamental roles. Most of the UPS and ALP genes were nevertheless differentially expressed across tissues, and their tissue-specific upregulation was associated with tissue-specific functions, phenotypes, and disease susceptibility. The small subset of UPS and ALP genes that was stably expressed across tissues was more highly and widely expressed and more essential for growth than other UPS and ALP genes, suggesting that it acts as a core. Lastly, we compared protein clearance to other branches of the proteostasis network. Protein clearance and folding were closely coordinated across tissues, yet both were less pivotal than protein synthesis. Taken together, we propose that the proteostasis network is organized hierarchically and is tailored to the proteome needs. This organization could contribute to and illuminate tissue-selective phenotypes.

## INTRODUCTION

Proteins in the cell are continuously synthesized and degraded, resulting in a highly dynamic proteome. The interplay between protein synthesis and clearance allows cells to replace existing proteins and adjust the proteome in response to internal or external cues. For example, regulatory or metabolic proteins, such as transcription factors or enzymes, are synthesized or degraded to modulate cellular responses to signals ^1,2^. Yet, relative to protein synthesis, protein clearance has diverse roles: Proteins are degraded into peptides to present samples of the cellular proteome to the immune system ^3^. Damaged proteins are degraded to prevent the accumulation of protein aggregates or misfolded proteins in the cell ^4,5^. Finally, unrequired proteins are degraded to provide amino acids, particularly in limiting conditions ^6^. Protein clearance in eukaryotic cells is carried concurrently by two major and highly conserved proteolytic systems, the ubiquitin-proteasome system (UPS) and the autophagy-lysosome pathway (ALP) ^1,7^.

The UPS is a selective proteolytic system that facilitates the degradation of a broad range of substrates by the proteasome through targeted ubiquitin conjugation ^8–11^. A cascade of three enzymes catalyzes the conjugation of ubiquitin to target proteins. An E1-activating enzyme forms a transient link with ubiquitin in an ATP-dependent manner. The activated ubiquitin is then transferred by an E2 conjugating enzyme to a substrate-bound E3 ligase. The E3 ligase, in turn, mediates the covalent attachment of ubiquitin to the substrate ^9,12^. Successive rounds of ubiquitylation assemble a polyubiquitin chain on the substrate, which can be trimmed by deubiquitinases (DUBs) ^9,13–15^. Similar conjugation cascades activate other type I ubiquitin-like proteins (UBLs), such as SUMO1-4 and UFM1, that, among other functions, promote or inhibit protein degradation ^16,17^. The UPS and most of its substrates are cytoplasmic, yet dedicated E3s and DUBs, aided by P97/VCP, enable protein retrotranslocation from the endoplasmic reticulum, mitochondria, and Golgi ^18–22^.

Human cells contain a few E1 enzymes, several E2 enzymes, and several hundreds of different E3 ligases and DUBs, allowing the UPS to process a broad range of substrates in parallel ^9^. The endpoint of the pathway, the 26S proteasome complex consisting of tens of proteins, is composed of a regulatory particle (19S) that specifically recognizes and unfolds ubiquitinated substrates and a core particle (20S) that processivity degrades them ^11,23,24^.

The ALP is an intracellular self-digestion pathway that ferries cytoplasmic constituents, such as dysfunctional, obsolete, or surplus proteins and organelles, to the lysosome, the endpoint of the pathway, for degradation and recycling ^25,26^. The cascade of macroautophagy (herein autophagy) is initiated by the UNC-51-like autophagy activating kinase (ULK) complex that activates the class III phosphatidylinositol 3-kinase complex I (PI3KC3-C1), generating phosphatidylinositol 3-phosphate (PI(3)P) and inducing phagophore nucleation ^27^. PI3P-binding proteins and lipids are then recruited to the elongating phagophore and, in turn, induce the lipidation of ATG8 orthologs (LC3 lipidation) by a ubiquitylation-like cascade ^26,27^. The maturation of the phagophore into the double membrane autophagosome is concluded by the endosomal sorting complexes required for transport (ESCRT) machinery that coordinates the closure of the autophagosome membrane ^27^. The autophagosome then fuses with the lysosome, resulting in cargo degradation ^28^. Cellular inputs, such as nutrient levels, stress signals, hormones, and available energy, can control the flux through the ALP by signaling pathways regulating autophagosome biogenesis and transcriptional and translational factors that modulate the expression of ALP genes ^25,29–31^.

The ALP encompasses nonselective and selective mechanisms of cargo recognition that converge on the lysosome for the degradation of their cargo ^26,32,33^. In nonselective autophagy, bulk cargo, such as portions of the cytosol, are engulfed within autophagosomes to recycle nutrients ^25^. Similarly, in general microautophagy, the lysosome takes up small amounts of cytoplasmic cargo via lysosomal membrane invaginations and engulfment ^28,34^. Whereas, in selective autophagy, LC3 interacts with cargo receptors that recognize specific proteins (often through ubiquitin modifications), selectively recruits the autophagy initiation machinery, and activates autophagosome biogenesis ^26,27^. For example, defective or surplus mitochondria can be selectively targeted for elimination by mitophagy receptors, such as OPTN and BNIP3 ^26^. In addition, both chaperone-assisted selective autophagy (CASA) and chaperone-mediated autophagy (CMA), jointly named chaperone-directed autophagy (CDA) ^35,36^, employ chaperones to select protein cargo for degradation ^37–39^.

Although protein clearance is fundamental to every cell, the demands for protein quality control could vary across tissues, cell types, and age ^40,41^. In mice, proteasome mapping revealed a tissue-specific sensitivity to protein aggregation that differed between males and females ^42^. Likewise, induced cranial and spinal motor neurons responded differently to protein misfolding, leading to distinct stress resistance ^43^. In humans, aortas transcriptomes revealed cell-type selective age-dependent changes in classes of the ALP ^44^. These varied demands for protein quality control also have pathological consequences. For example, certain mutations in the E3 ligase Parkin and the ubiquitinated protein shuttle, UBQLN2, increase the risk for late-onset Parkinson’s disease and Amyotrophic lateral sclerosis, respectively ^45^. Tissue contexts also affect the crosstalk between UPS and ALP. In *C. elegans*, compromised autophagy affected UPS function in a tissue-specific manner ^46^. Likewise, in aging neurons, a switch between the relative expression levels of BAG1 and BAG3 shifted the degradation of protein aggregates from UPS to ALP ^47^. We recently studied the organization of the chaperone system, a different branch of the proteostasis network ^48^. We found that the chaperone system is composed of genes with tissue-selective expression and functionalities, and more essential genes with steady expression across tissues and throughout development. Yet, the organization of the human clearance systems and subsystems across tissues and cell types under physiological conditions remains elusive.

Here, we interrogated the basal organization of UPS and ALP across human tissues and cell types. We found that relative to protein-coding genes, UPS and ALP were more widely and uniformly expressed across adult tissues and cell types, and during development. They declined more with age and were more essential for growth. Nevertheless, classes of UPS and ALP, such as the proteasome and the lysosome, behaved variably with respect to expression patterns, essentiality, and disease susceptibility. Most UPS and ALP genes were differentially expressed across tissues, and expression upregulation was associated with tissue-selective functions, as demonstrated using *C. elegans*, and disease manifestation. Similarly to the chaperone system, UPS and ALP contained a small core subset of more essential genes with wider expression across tissues, cell types, and age. Lastly, we compared protein clearance and other proteostasis branches, including protein synthesis and protein folding. Protein folding was closely coordinated with UPS and ALP, while protein synthesis was less coordinated and more fundamental, as demonstrated by its essentiality and wider expression. The plasticity of the proteostasis network is linked to tissue-specific proteostasis requirements and can contribute to tissue-selective protein misfolding diseases, such as neurodegenerative disorders and myopathies.

## RESULTS

### UPS and ALP are fundamental across tissues

To examine the behavior of UPS and ALP across tissues and cell types, we started by defining the components of each system. For that, we used the comprehensive enumerations of the different branches of the human proteostasis system, including gene classes, groups, and types, that were recently compiled by the proteostasis consortium ^35,36,49^. Accordingly, UPS included E1 activating enzymes, E2 conjugating enzymes, E3 ubiquitin ligases, p97 and associated proteins, ubiquitin- and ubiquitin-like-proteins (‘Ub and UBL’), and proteasome and associated proteins, summing up to 1188 genes (see Methods, Table S1A). ALP included autophagy preinitiation signaling, gene expression, substrate selection, CDA, initiation and elongation, autophagosome closure maturation, lysosome maturation reformation and fusion, ESCRT, and lysosome catabolism, summing up to 829 genes (see Methods, Table S1B). Certain genes belonged to multiple classes, such as ATXN3 and ATG12, and few, such as VCP, were shared between the two clearance systems.

We first analyzed the expression patterns of UPS and ALP genes across human tissues. For this, we used transcriptomes of 34 human tissues sequenced in bulk ^50^ and transcriptomes of 28 human tissues sequenced using single-cell RNA sequencing ^51^. Per dataset, we associated each gene with the fraction of tissues expressing it in bulk or the fraction of cell types per tissue expressing it (see Methods). According to each of these datasets, UPS and ALP genes were more ubiquitously expressed than other protein-coding genes (Mann-Whitney (MW) test, adjusted *P*<1e-41; Fig. 1A-B, Fig. S1A-B). For example, using the bulk dataset, UPS and ALP genes were expressed in 95% of the tissues relative to 74% for other protein-coding genes (medians reported, Fig. 1A). To confirm the relevance of these observations to the proteome, we turned to a proteomic dataset of 32 human tissues ^52^. As explained therein, proteins were classified as tissue-specific (TS), tissue-enriched (TE), housekeeping, or other (see Methods). For simplicity, we combined the tissue-specific and tissue-enriched proteins into one group, denoted TS/TE. UPS and ALP systems included more housekeeping proteins and fewer TS/TE proteins compared to other proteins (MW test, *P*<2.6e-11 and *P*<0.005, respectively; Fig. 1C), in accordance with the transcriptomic datasets.

**Figure 1:**
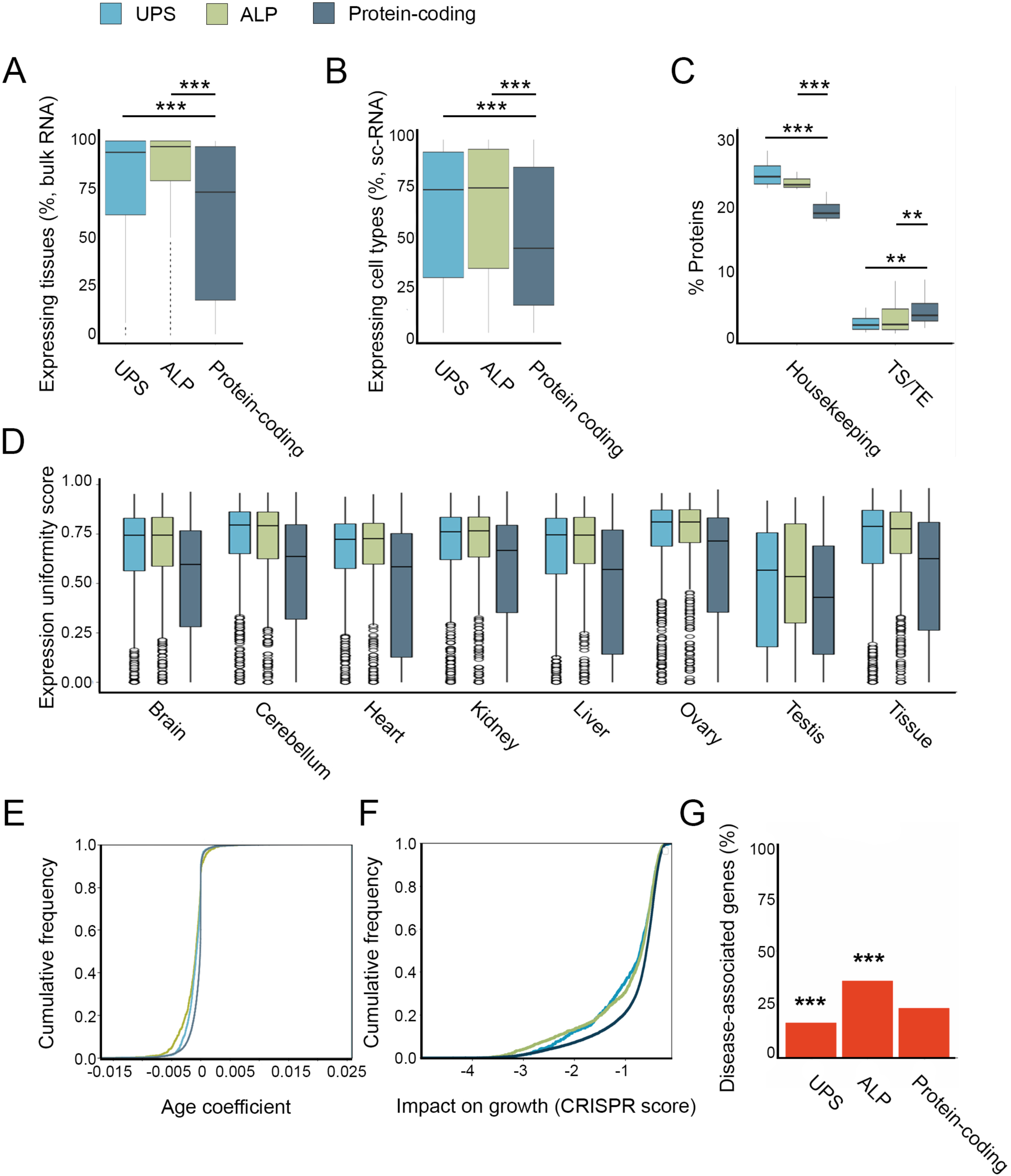
Expression patterns across tissues and time and phenotypic impacts distinguish UPS and ALP from other protein-coding genes. A. The percent of adult human tissues that express UPS, ALP, and other protein-coding genes. Analysis was based on bulk RNA transcriptomes of 34 tissues. Data included 1127 UPS, 812 ALP, and 16,771 other protein-coding genes that were expressed in at least one tissue. UPS and ALP genes were significantly more ubiquitous than other protein-coding genes (MW, adjusted *P*<9.6e-41 and *P*<2.3e-75). B. The percent of cell types per tissue that express UPS, ALP, and other protein-coding genes. Analysis was based on single-cell transcriptomes of 28 tissues. Data included 1078 UPS, 820 ALP, and 14,148 protein-coding genes that were expressed in at least one cell type. UPS and ALP genes were significantly more ubiquitous than other protein-coding genes (MW, adjusted *P*<e-100 in both). C. The fraction of housekeeping and tissue-specific/tissue-enriched (TS/TE) proteins per system, according to proteomic analysis of 32 adult human tissues. UPS and ALP were enriched for housekeeping proteins (MW, adjusted *P*<2.6e-11) and depleted of TS/TE proteins (MW, adjusted *P*< 0.005) relative to other protein-coding genes. Data included 809 UPS, 706 ALP, and 10,271 protein-coding genes. D. Expression uniformity per tissue across time and across tissues of UPS, ALP, and other protein-coding genes. Analysis was based on bulk transcriptomes of seven human tissues in development and age. UPS and ALP genes were more uniformly expressed than other protein-coding genes (MW, adjusted *P*=1.8e-60 and 1.3e-60). Data included 1132 UPS, 700 ALP, and 14,648 protein-coding genes. E. The age coefficient of mouse orthologs of UPS, ALP, and other protein-coding genes. Age coefficient reflects the change in transcript level with age across 23 mouse tissues, and each gene was associated with its median age coefficient. UPS and ALP had lower age coefficients than other protein-coding genes (KS test, adjusted *P*<8.2e-51 and 9.9e-52, respectively). Data included 999 UPS, 736 ALP, and 13,148 protein-coding genes. F. The impact on cell growth of UPS, ALP, and other protein-coding genes, measured using 1078 cell lines harboring CRISPR-induced gene inactivation (CRISPR score). Each gene was associated with its minimal CRISPR score. UPS and ALP were more essential for growth than other protein-coding genes (KS test, adjusted *P*<1.6e-13). Data included 1062 UPS, 673 ALP, and 13,717 protein-coding genes. G. The fraction of Mendelian disease genes per system, out of the total numbers of genes in Fig. 1A. UPS (187 genes) was depleted of Mendelian disease genes, and ALP (294 genes) was enriched for them (adjusted *P*=3.6e-9 and 2.4e-16, respectively; Fisher exact test), relative to other protein-coding genes (3919 genes).

The abovementioned transcriptomic and proteomic datasets were obtained by sampling adult donors. Given the deterioration of the proteostasis system with age ^29,53,54^, it was intriguing to assess the ubiquity of the two clearance systems over time. We first utilized bulk transcriptomic analysis of seven human organs sampled from embryonic stages to adulthood ^55^. Therein, each gene was scored per organ by the uniformity of its expression across time points. In all organs, UPS and ALP genes were more uniformly expressed than other protein-coding genes (MW test, adjusted *P*=1.8e-60; Fig. 1D). A similar tendency for uniformity was observed upon scoring genes by the uniformity of their expression across organs (MW test, adjusted *P*<2.5e-36, Fig. 1D). Next, we utilized transcriptomic analysis of 24 mouse tissues across ages, whereby the correlation between the expression level of each gene and age was calculated per tissue (see Methods). In accordance with the decline in functionality of the proteostasis system with age ^54^, both UPS and ALP genes were more negatively correlated with age than other protein-coding genes (Kolmogorov-Smirnov (KS) test, adjusted *P*<1e-50 for both; Figure 1E). Of note, the decline with age was more pronounced in ALP relative to UPS genes (KS test, *P*=0.005).

Lastly, we assessed the essentiality of each system, as determined by the impact of its genes on cell growth and their involvement in Mendelian diseases. For gene impact on cell growth, we harnessed data from the Cancer Dependency map (DepMap^56^), where the impact of CRISPR gene knockouts on cell growth was measured per gene in 1078 cell lines (see Methods). UPS and ALP genes were more essential than other protein-coding genes (KS test, *P*=1.6e-13, and *P*=9.2e-14, respectively; Fig 1F). For Mendelian diseases, we mined OMIM for genes whose aberration was causally associated with Mendelian diseases (i.e., disease genes, see Methods). In total, 187 UPS genes (17%) and 294 ALP genes (36%) were associated with Mendelian diseases (Table S2). Given that 31% of the other protein-coding genes were associated with Mendelian diseases, UPS was depleted of them, whereas ALP was enriched for them (adjusted *P*=3.6e-9 and *P*=2.4e-16, respectively, Fisher exact test; Fig. 1G). Taken together, relative to other genes and proteins, UPS and ALP components were more widely and uniformly expressed across tissues, cell types, and age, declined more with age, and were more essential for growth, in accordance with their fundamental roles in living cells.

### UPS and ALP classes show distinct behaviors

We next explored the organization of each clearance system at a finer resolution, starting with the UPS (Fig. 2A, Table S1A). We examined the expression pattern of the different UPS classes across tissues using the bulk and single-cell transcriptomic and proteomic datasets. In accordance with Fig. 1A-C, the different classes were highly ubiquitous across tissues and cell types (Fig. 2B-D). Yet, the proteasome and associates (henceforth denoted proteasome, 82 genes) was the most ubiquitous class, whereas E3, the largest class (706 genes), was the most varied across tissues (MW test: bulk p<e-10; single-cell: *P*<e-100). Lastly, we examined the essentiality of each class with respect to the impact of its genes on cell growth and involvement in Mendelian diseases (Fig 2E-F). E1 and the proteasome were the most essential (MW test, *P*=0.0009 and *P*=1.3e-12, respectively), whereas E3 was the least essential (MW test, *P*=1.6e-9). E3 also contained the largest fraction of Mendelian disease genes (115 genes, 62%; Fig. 2F). One example is the E3 ligases, KLHL40 and KLHL41, that are causal for Nemaline myopathy. The proteasome class contained nine Mendelian disease genes, five of which are causal for the same disease, proteasome-associated autoinflammatory syndromes (PRAAS)^57^. These included the three specialized catalytic subunits of the immunoproteasome, PSMB8, PSMB9, and PSMB10, the constitutive PSMB4 subunit, and the proteasome maturation protein, POMP, that preferentially promotes immunoproteasome assembly ^58,59^. Taken together, UPS classes show differential impacts on growth and disease.

**Figure 2:**
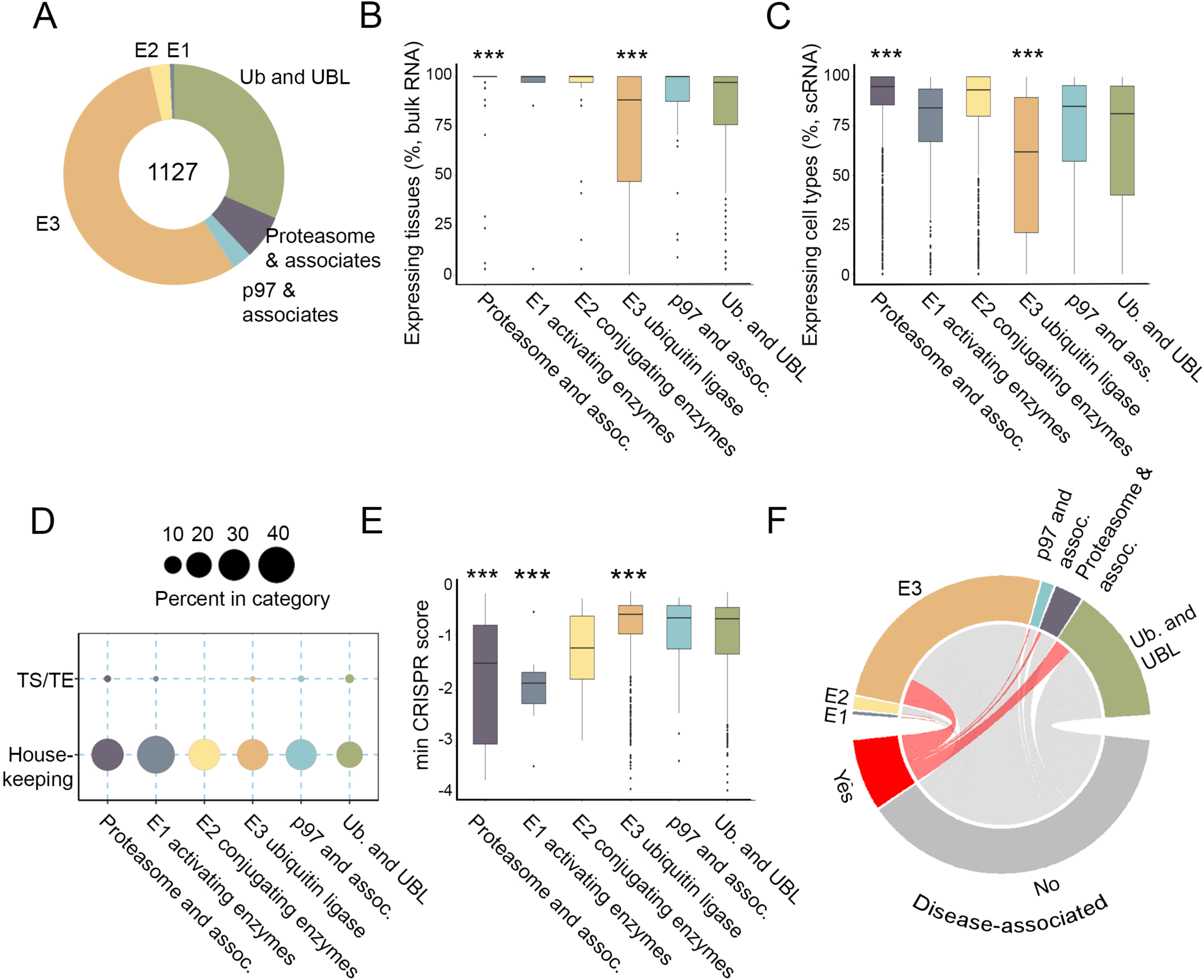
Variability in expression patterns and phenotypic impact among UPS classes. A. The relative number of genes per UPS class out of a total of 1127 genes. The number of genes per class: proteasome and associates, 82; E1, 9; E2, 36; E3, 706; p97 and associates 36; Ub and UBL, 400. Genes may belong to multiple classes. B. The percent of adult human tissues that express UPS genes per class. Proteasome and associates class was expressed in all tissues, whereas E3 was the most diverse (adjusted *P*=2.1e-11 and 2.4e-16, respectively; MW). C. The percent of cell types per tissue that express UPS genes per class. Proteasome and associates class was the most ubiquitous, whereas E3 was the most specific (adjusted *P*=7.5e-297 and e-100, respectively; MW). D. The fraction of housekeeping and TS/TE proteins per UPS class. E1 had the highest fraction of housekeeping proteins. E. The impact on cell growth of UPS classes. E1 and Proteasome had the highest impact on growth, whereas E3 had the lowest impact (adjusted *P*=8.9e-4, 1.3e-12, and 1.6e-9, respectively; MW). F. The fraction of Mendelian disease genes per class.

To ask whether a highly ubiquitous class could show variability, we focused on the proteasome and its associated groups, including the proteasome core subunit (20S), regulatory subunit (19S), adaptors, activators and inhibitors, assembly chaperones, and other genes (Fig. S2A). All groups were highly ubiquitous across tissues and cell types, with the exception of assembly chaperones (Fig. S2B-C), and contained large fractions of housekeeping proteins (Fig. S2D). In contrast to this uniformity, the essentiality of proteasome subgroups was highly variable. The proteasome core and regulatory subunits were significantly more essential than other subgroups (MW test, *P*=0.0005, Fig. S2E). Yet, the specialized core subunits, including the testis-specific proteasomal subunit PSMA8 required for the spermatoproteasome ^60^, and the three immunoproteasomes subunits PSMB8, PSMB9, and PSMB10, linked to PRAAS disease ^58,59^, were not essential (Fig. S2E-F).

We conducted similar analyses of ALP classes (Fig. 3A, Table S1B). Three classes with interesting behaviors were ESCRT, CDA, and lysosome catabolism. ESCRT and CDA genes were among the most ubiquitous classes across tissues and cell types (Fig. 3B-C; MW test: bulk *P*≤0.09; single-cell: *P*≤8.2e-13) and included the largest fraction of housekeeping genes (52% and 49%, Fig. 3D). Whereas ESCRT was the most essential class (MW test, adjusted *P*=0.0002, Fig. 3E) and was relatively depleted of disease genes (19%, Fisher exact test, adjusted *P*=0.055, Fig. 3F), CDA was less essential (MW test, adjusted *P*=0.043, Fig. 3E). Lysosome catabolism genes, in contrast, were more variable than other classes across tissues and cell types (Fig. 3B-C) and included the smallest fraction of housekeeping genes (15%, Fig. 3D). They were also less essential (MW test, *P*=1.4e-6; Fig. 3E), and were enriched for Mendelian disease genes (48%, Fisher exact test, *P*=0.004, Fig. 3F). Taken together, while the UPS and ALP appear to be fundamental, their classes behave variably with respect to expression patterns, essentiality, and disease susceptibility.

**Figure 3:**
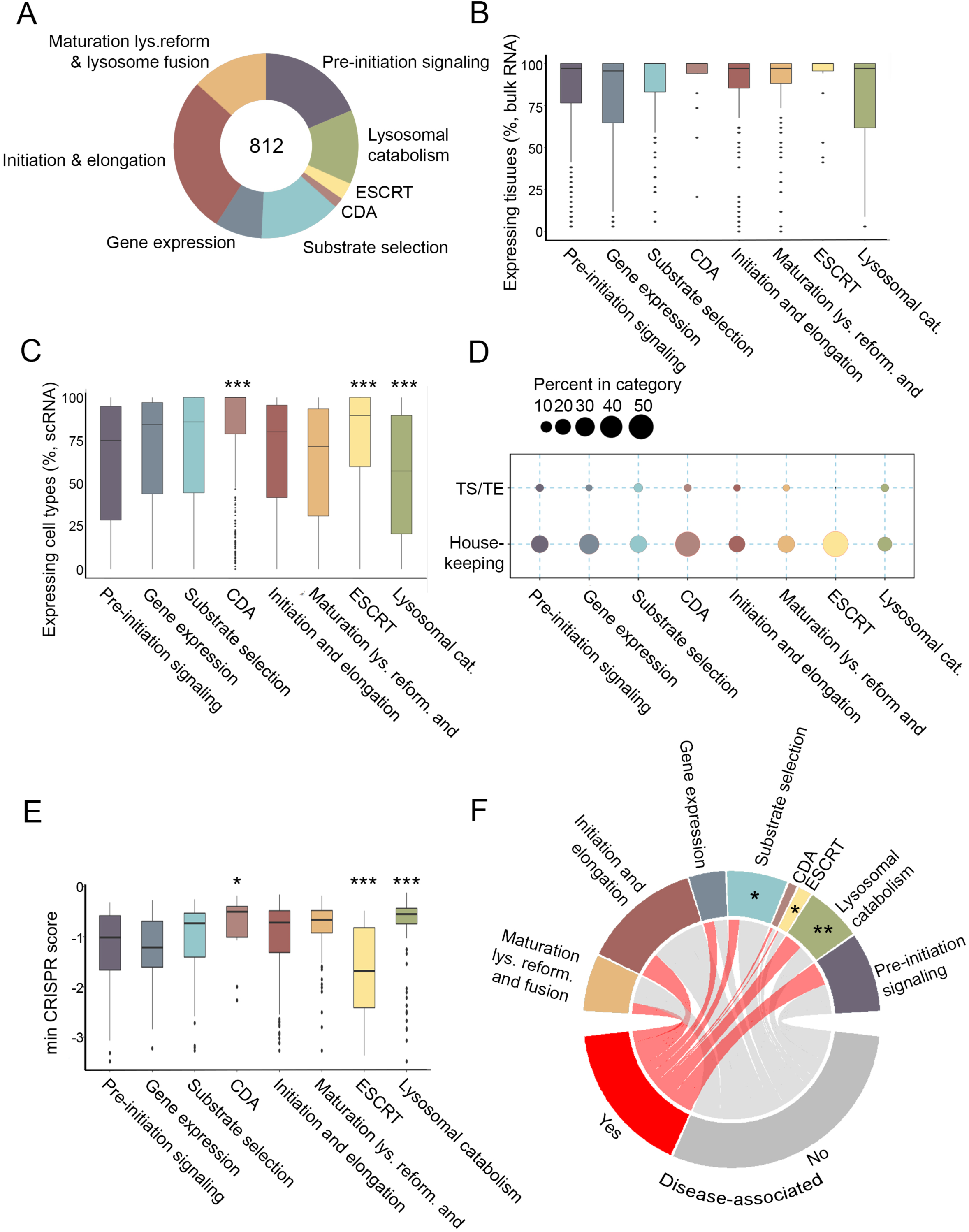
Variability in expression patterns and phenotypic impact among ALP classes. A. The relative number of genes per ALP class out of a total of 812 genes. The number of genes per class: Pre-initiation signaling, 167; gene expression, 74; substrate selection, 127; chaperone-directed autophagy (CDA), 17; initiation and elongation, 247; maturation, reformation and lysosome fusion, 115; ESCRT, 27; lysosomal catabolism, 116. Genes may belong to multiple classes. B. The percent of adult human tissues that express ALP genes per class. ESCRT and CDA were the most ubiquitous (adjusted *P*=0.08 and 0.09, respectively; MW). C. The percent of cell types per tissue that express ALP genes per class. ESCRT and CDA were the most ubiquitous, whereas lysosome catabolism was the most specific (adjusted *P*=8.2e-13,1.3e-33, and 5.8e-69, respectively; MW). D. The fraction of housekeeping and TS/TE proteins per ALP class. ESCRT had the highest fraction of housekeeping proteins and the lowest fraction of TS/TE proteins. E. The impact on cell growth of ALP classes. ESCRT had the highest impact on growth, CDA was less essential, whereas lysosome catabolism had the lowest impact (adjusted *P*=0.0002, 0.043, and 1.35e-6, respectively; MW). F. The fraction of Mendelian disease genes per ALP class. Substrate selection and ESCRT were depleted of disease genes, whereas lysosome catabolism was enriched for them (adjusted *P*=0.016, 0.055, and 0.004, respectively; Fisher exact test).

### Most UPS and ALP genes are differentially expressed across tissues

We previously showed that the chaperone system consists of a majority of genes with tissue-selective expression and functions and a small subset of core genes with stable expression and essential functions ^48^. Given that chaperones are part of the proteostasis system and share similar fundamental features as UPS and ALP (Fig. 1), we next examined whether they also share a similar organization. To test for tissue-selective expression of UPS and ALP genes, we performed a differential expression analysis, comparing the bulk transcriptome of each tissue against all other tissues (see Methods). Both systems displayed tissue-sensitive expression patterns, with most genes showing over 2-fold upregulation or downregulation in at least one tissue (Fig. 4A). Skeletal muscle, whole blood, testis, and brain tissues showed the highest fraction of differentially expressed UPS and ALP genes (Fig. 4A). Notably, manual curation of UPS and ALP Mendelian disease genes showed that these tissues were also the most susceptible to tissue-selective Mendelian diseases, including neurological disorders, blood disorders, and myopathies (Fig. 4B and Table S2). For example, the topmost UPS and ALP upregulated genes in skeletal muscle were the Nemaline myopathy genes KLHL40 and KLHL41 E3 ligases and PRKAG3 gene linked to skeletal muscle excess glycogen, respectively ^61–64^. These data suggest that the upregulation of UPS and ALP genes is functionally relevant and linked to tissue-specific phenotypes.

**Figure 4:**
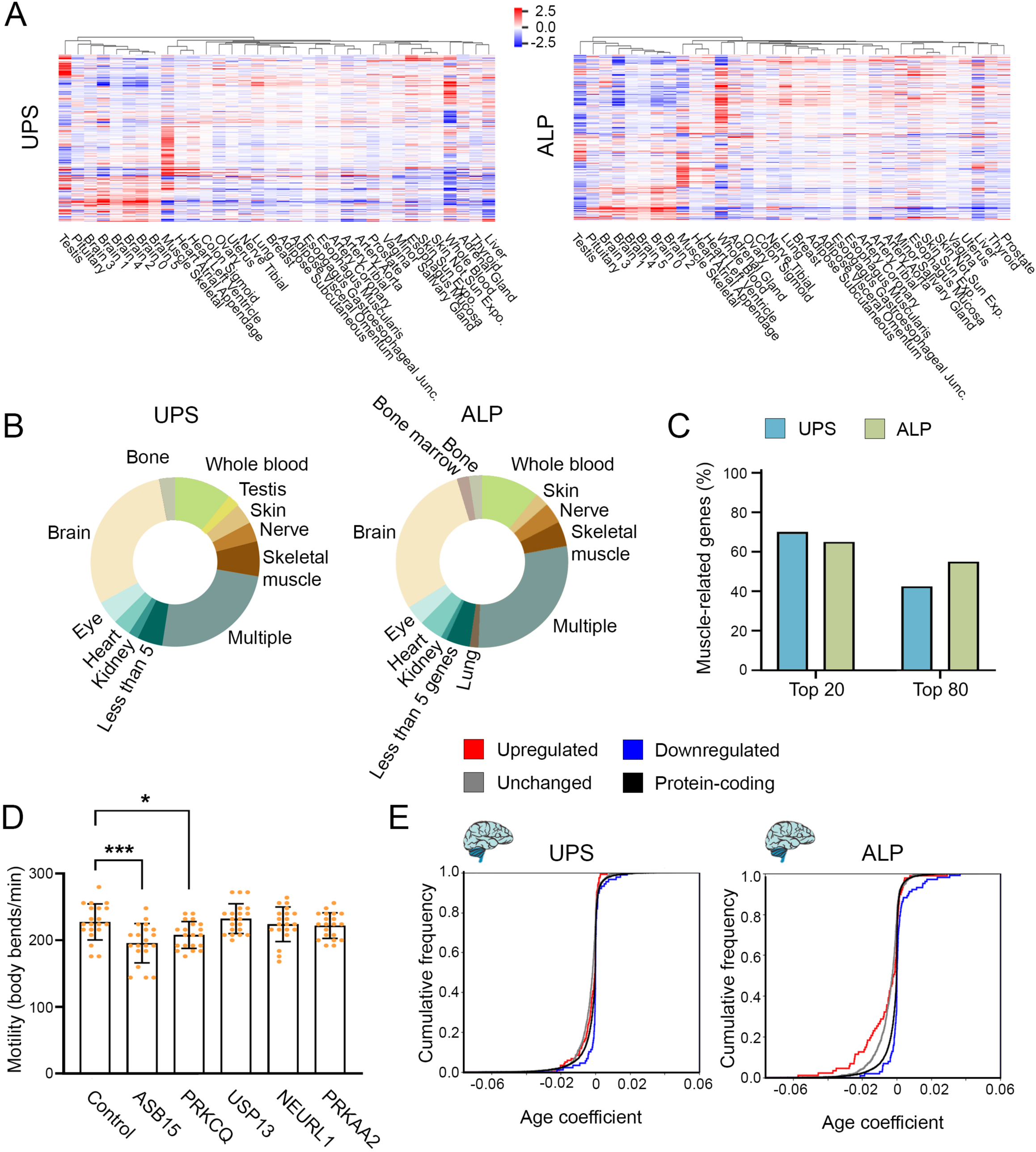
UPS and ALP genes show tissue-dependent expression and impacts. A. The differential expression of UPS (left) and ALP (right) genes in adult human tissues. Both systems display tissue-specific expression patterns. B. The fraction of Mendelian disease genes with tissue-specific clinical phenotypes per system. The largest number of disease genes affect the brain or multiple tissues. Other corresponds to tissues with less than five Mendelian disease genes. C. The fraction of skeletal muscle-upregulated genes with muscle-linked phenotypes per system. The percent of UPS or ALP skeletal muscle-upregulated genes (out of the top 20 or 80 upregulated genes) that were associated with muscle phenotype based on manual literature curation. D. Reduced motility of RNAi-treated *C. elegans*. The expression of worm orthologs of UPS and ALP skeletal muscle-upregulated genes (five genes not previously associated with muscle function) was knockdown using RNAi and worms’ motility was scored (n=20). Worms treated with RNAi against ASB15 and PRKCQ orthologs, *M60.7* and *B0545.1* respectively, resulted in reduced motility. Data are means ± 1 standard error of the mean (1SE). Data were analyzed using one-way ANOVA followed by a Dunnett’s post-hoc test. (*) denotes *P*≤0.05, (**) denotes *P*≤0.005 compared with empty vector control. E. The age coefficient in non-myeloid brain tissue of mouse orthologs of UPS and ALP genes. UPS genes that were upregulated (118 genes), unchanged (784 genes), or downregulated (88 genes) in human brain tissue had similar age coefficients, except for downregulated genes that had higher age coefficients relative to unchanged genes (KS test, adjusted *P*<4.9e-14). ALP genes that were upregulated (87 genes) or unchanged (543 genes) in human brain tissue declined more rapidly with age than downregulated (101 genes) genes or other protein-coding genes (KS test, adjusted *P*<4.3e-6 and 2.5e-16, respectively).

Next, we tested whether tissue-specific upregulation could have functional implications. We focused on the top 80 upregulated genes in skeletal muscle per system. We first checked whether these genes were associated with muscle phenotypes, using manual literature curation (see Methods). We associated 43% and 55% of the UPS and ALP genes, respectively, with muscle phenotypes (Table S3). These percentages were even higher among the top 20 muscle-upregulated genes (70% and 65%, respectively; Fig. 4C). Lastly, we experimentally tested the association of muscle-upregulated genes with muscle function using *Caenorhabditis elegans*. We focused on five of the top UPS and ALP muscle-upregulated genes that were not previously associated with muscle function and have orthologs in worms (see Methods). We then examined whether expression knockdown of these five genes will impact animals’ motility. Of the genes tested, RNAi knockdown of two genes, ASB15 (UPS) and PRKCQ (ALP), resulted in reduced motility rates compared to the control (86±3% and 91±2%; Fig. 4D), suggesting that they have a role in muscle function. Taken together, the UPS and ALP muscle-selective expression patterns are functionally relevant.

We next asked whether the upregulated and downregulated genes of a given tissue had distinct correlations with age. We focused on brain and skeletal muscle tissues that are linked to many age-dependent diseases in which proteostasis decline is implicated ^4,29,65–68^. In brain, ALP genes that were upregulated or unchanged declined more rapidly with age than downregulated genes or other protein-coding genes (Fig. 4E). UPS, in contrast, did not show such changes. In skeletal muscle, no differences were observed (Fig. S3). This suggests that the brain-active components of ALP decline with age in a tissue-dependent manner. In agreement, aged brain-associated decline in autophagy gene expression and function was observed in worms, mice, and humans, and several neurodegenerative diseases are linked to age-dependent deficits in autophagy ^29,41^.

### The UPS and ALP clearance systems contain a small set of core genes

Differential expression analysis revealed that both UPS and ALP contain a small subset of genes that were stably expressed in all tissues (see Methods; UPS: 111/1,127, 10%; ALP 64/812, 8%). For example, UBA1, the main E1-ubiquitin-activating enzyme ^69^, and ATG13, part of the ULK complex and a target of several upstream signaling pathways that regulate autophagosome biogenesis ^27,70^. These core subsets included representatives from each class yet were not uniformly distributed (Fig. 5A). In UPS, this subset was significantly enriched among E2s and relatively depleted from E3s (adjusted *P*=0.005 and *P*=0.0026, respectively, one-sided Fisher exact test). In ALP, this subset was relatively enriched among ESCRT and relatively depleted from lysosome catabolism genes (adjusted *P*=0.1 and *P*=0.0018, respectively, one-sided Fisher exact test).

**Figure 5:**
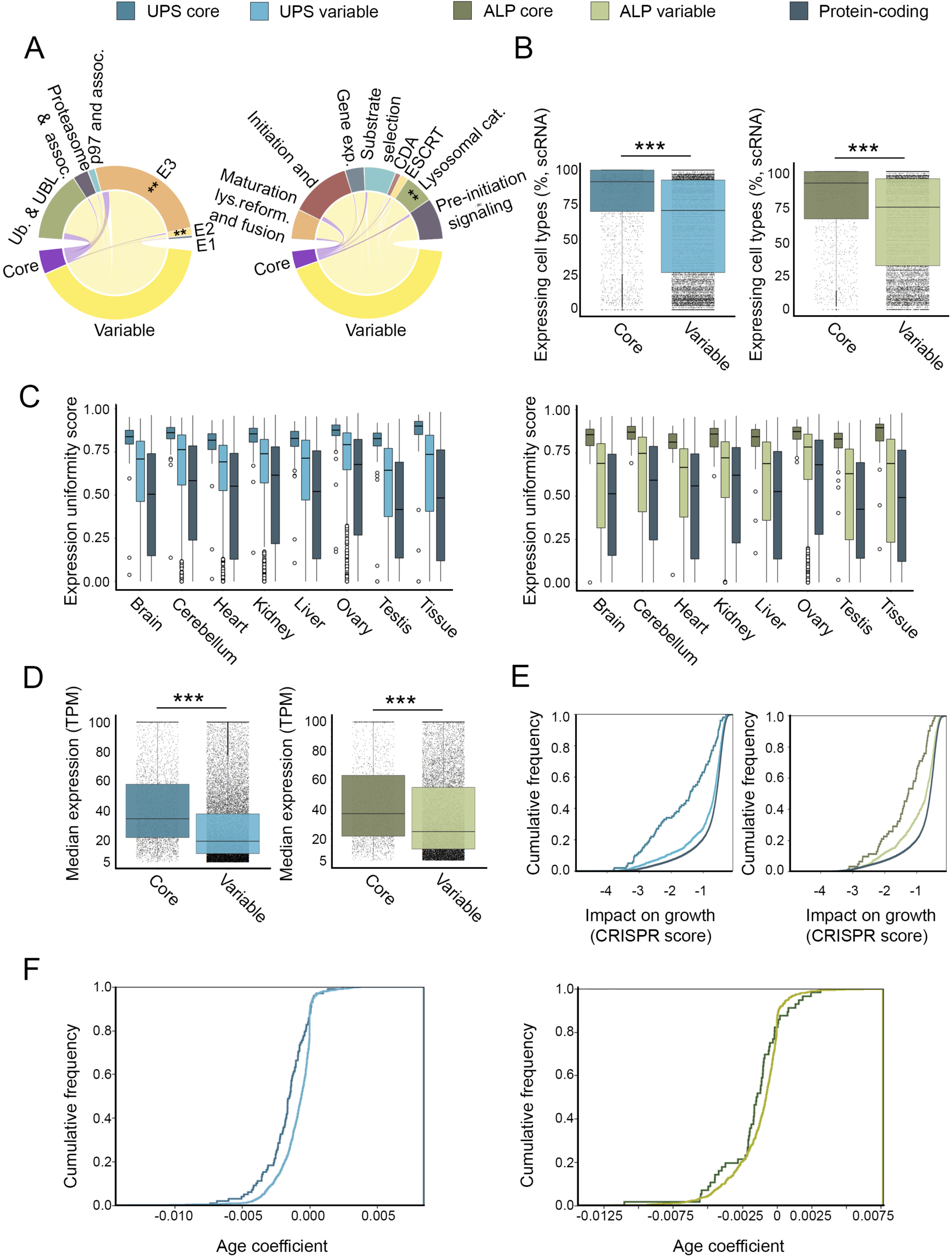
Stably expressed genes within UPS and ALP behave as core subsystems. A. The fraction of stably and variably expressed genes in UPS (left) and ALP (right). In UPS, stably expressed genes were enriched among E2 and relatively depleted from E3 (adjusted *P*=0.005 and 0.0026, respectively, one-sided Fisher exact test). In ALP, they were enriched among ESCRT and relatively depleted from lysosome catabolism (adjusted *P*=0.1 and 0.0018, respectively, one sided-Fisher exact test). B. The percent of cell types per tissue expressing UPS (left) and ALP (right) genes defined in A as stably or variably expressed, per system. Stably expressed genes were more broadly expressed across cell types (MW test, p=2.5e-159 and p=1.1e-61, respectively). C. Expression uniformity across time and tissues of stably and variably expressed UPS (left) and ALP (right) genes per system. Stably expressed genes were expressed more uniformly across time and tissues (MW test, *P*<2.6e-36 and 8.8e-16, respectively). D. Median expression levels in adult human tissues of stably and variably expressed UPS (left) and ALP (right) genes per system. Stably expressed genes were more highly expressed (MW test, *P*=3.2e-284 and 2.3e-65, respectively). E. The impact on cell growth of stably and variably expressed UPS (left) and ALP (right) genes per system. Stably expressed genes had a larger impact on cell growth (KS test, *P*<1.7e-11 and 2.4e-6, respectively). F. The age coefficient of mouse orthologs of stably and variably expressed UPS (left) and ALP (right) genes per system. Stably expressed genes declined more with age (KS test, *P*=2.3E-7 and 0.003, respectively).

To test the ‘coreness’ of these stably expressed subsets, we first tested whether they were also stably expressed in cell types and across age. Indeed, stably expressed genes were expressed in more cell types than other UPS and ALP genes (MW test, *P*=2.5e-159 and *P*=1.1e-61, respectively, Fig. 5B), and were expressed more consistently than other genes across organs and time points (MW test, *P*<2.6e-36 and *P*<8.8e-16, respectively; Fig. 5C). We also compared their bulk expression levels across tissues to the levels of the remaining UPS and ALP genes. In both clearance systems, the stably expressed subsets were more highly expressed than the remaining genes (MW test, *P*=3.2e-284 and *P*=2.3e-65, respectively; Fig 5D).

Next, we assessed the essentiality of the stably expressed subset of each system. We found them to be significantly more essential than other UPS and ALP genes (KS test, *P*<1.7e-11 and *P*<2.4e-6, respectively; Fig. 5E). When examining disease-association, they were not significantly enriched or depleted of Mendelian disease genes (Table S2). Interestingly, even a synonymous mutation in the stably expressed E1 gene, UBA1, led to spinal muscular atrophy, suggesting that small changes in its expression could lead to a severe phenotype ^71^. Lastly, we tested whether stably expressed genes had different correlations with age than other UPS and ALP genes. In UPS and slightly less so in ALP, stably expressed genes declined more with age (KS test, *P*=2.3E-07 and *P*=0.003, Fig. 5F). Taken together, the stably expressed genes of each clearance system emerge as core components of UPS and ALP across tissues, cell types, and age.

### Coordinated variability of UPS and ALP genes across tissues

Both UPS and ALP are responsible for protein clearance. Given that many of their genes had varied expression in distinct tissues, we asked whether their expression changed in a coordinated or complementary manner. For that, we computed the fractions of upregulated and downregulated genes (>2-fold change) in each tissue and compared them between clearance systems (Fig. 6A). We found that the upregulation and downregulation of UPS genes was correlated with the up- and downregulation of ALP genes (Pearson r=0.86 and r=0.89, respectively). A similar tendency was shown for the fraction of TS/TE proteins (Pearson r=0.91, Fig. 6B). Since cancer is associated with disruption of cellular signaling and regulation, we asked whether the coordinated expression of UPS and ALP is maintained in cancer tissues. For this, we repeated our analyses using cancer transcriptomic data from the Cancer Genome Atlas (TCGA, see Methods). Like normal tissues, the correlation patterns between the different clearance systems were maintained in cancer (Pearson r=0.84 and r=0.95, respectively, Fig. 6C). These data suggest that the clearance systems are highly coordinated.

**Figure 6.**
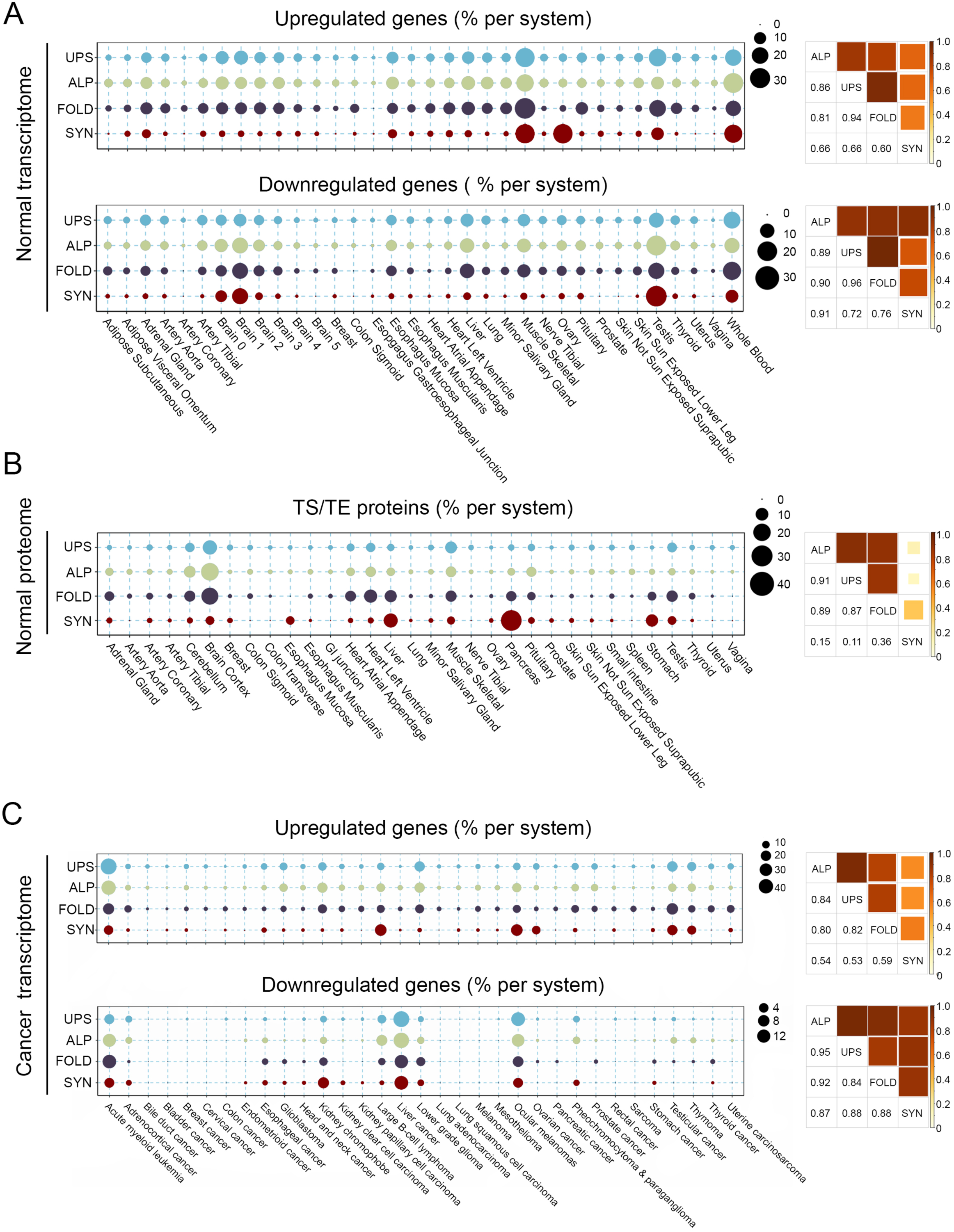
Proteostasis branches are correlated in their expression across tissues and cancers. Proteostasis branches included UPS, ALP, protein folding (FOLD), and protein synthesis (SYN). Spearman correlations between branches were computed. A. The fractions of upregulated (top) and downregulated (bottom) genes per system in adult human tissues (expression fold-change >=2 and <=0.5, respectively). B. Fractions of TS/TE proteins per system in adult human tissues. C. The fractions of upregulated (top) and downregulated (bottom) genes per system in human cancers (preferential expression >=2 and <=0.5, respectively).

We previously showed that UPS and ALP classes are not similarly expressed across tissues (Figs. 2-3). We next asked whether these classes were upregulated in a coordinated manner within a tissue. We focused on the tissues where gene upregulation was most frequent, including skeletal muscle, whole blood, testis, and brain tissues (Fig. 4A, 6A). We found that classes differed within a tissue and between tissues (Fig. S4A). We further analyzed the coordination among groups of the proteasome class. In brain tissue, proteasome groups were mostly unchanged. However, in skeletal muscle tissue, all proteasomal groups were upregulated (Fig. S4B). In particular, 16/19 (84%) of the proteasome regulatory genes were upregulated in skeletal muscle tissue versus none in brain tissue (Fig. S4C) in agreement with tissue-specific differences in proteasome activity, stress sensitivity, and disease susceptibility ^42,72^. Altogether, UPS and ALP were coordinated at the system level, but classes and their groups had varied and tissue-selective behaviors.

### The relationships between protein clearance, synthesis, and folding systems across tissues

The coordinated expression of the UPS and ALP clearance systems could be driven by tissue-specific requirements and protein-misfolding loads. We therefore examined the behavior of protein synthesis and protein folding systems across tissues, composed of 324 and 294 genes, respectively (see Methods). We first examined the expression patterns of these systems across tissues. The protein folding system was variably expressed across tissues and correlated with the two clearance systems at both the gene and protein levels (Pearson r≥0.81 and Pearson r≥0.87, respectively, Fig. 6A-B). A similar pattern of coordination between the different systems was also observed across cancers (Fig. 6C). In contrast, the expression of the protein synthesis system was relatively stable across tissues and accordingly less correlated with the two clearance systems (Pearson r≤0.66. and r≤0.15, respectively, Fig. 6A-B). Still, protein synthesis was upregulated in skeletal muscle and whole blood tissues, and uniquely upregulated in ovary. The upregulation of protein synthesis in skeletal muscle and whole blood tissues could explain the upregulation of the folding and clearance systems in those tissues. However, it is not the main driver of protein folding and clearance upregulation in other tissues.

The less variable expression of the protein synthesis system led us to compare the behaviors of the different systems, including expression prevalence in tissues and cell types and essentiality. We found that protein synthesis was indeed more ubiquitously expressed across tissues and cell types (Fig. 7A-B) and contained a higher fraction of house-keeping proteins (Fig. 7C). It was also more uniform across age (Fig. 7D). Although the protein synthesis system was smaller in size, its median expression was significantly higher (MW test, p≤1.4e-6, Fig. 7E). Protein synthesis was significantly more essential for growth (KS test *P*≤1.1e-26, Fig. 7F), but had a similar fraction of Mendelian disease genes (Fig. 7G). Interestingly, the decline of protein synthesis with age in mice was less severe than in other proteostasis systems, and some genes even increased with age (Fig. 7H). Taken together, protein synthesis appears to be the most fundamental among the proteostasis systems, followed by protein folding and the two protein clearance systems.

**Figure 7.**
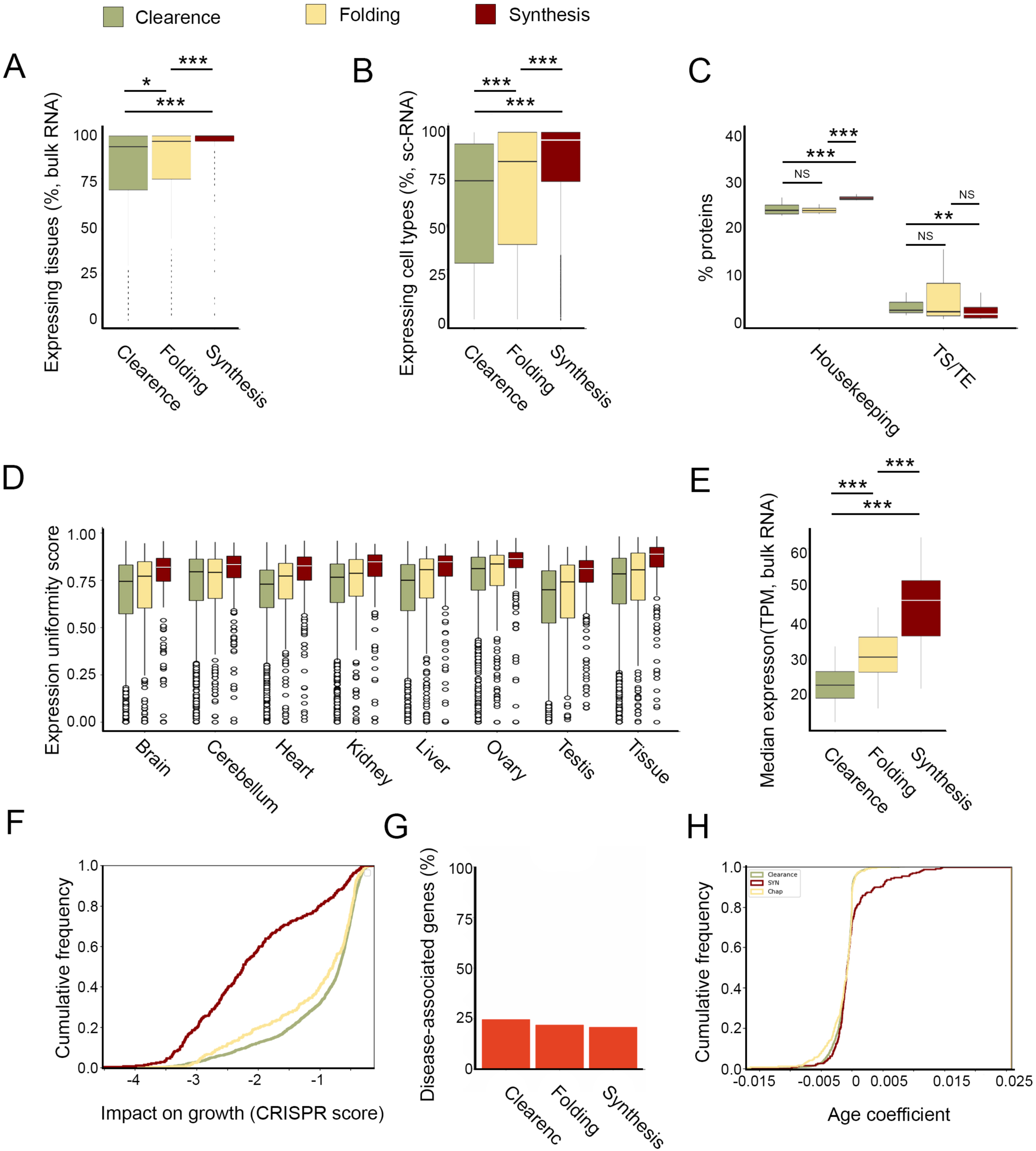
Expression patterns across tissues and time and phenotypic impacts of proteostasis systems. For simplicity, UPS and ALP were combined into a single clearance system. A. The percent of adult human tissues that express each proteostasis system. Protein synthesis was most widely expressed, followed by protein folding (adjusted *P* synthesis vs. clearance:1.3e-26; folding vs synthesis: 1.3e-11, folding vs. clearance: 0.02; MW). B. The percent of cell types per tissue that express each system. Protein synthesis was most widely expressed, followed by protein folding (adjusted *P* synthesis vs. clearance:<e-100; folding vs synthesis: 6.1e-172, folding vs. clearance: 1.6e-54; MW). C. The fraction of housekeeping and TS/TE proteins per system. Protein synthesis contained a higher fraction of housekeeping proteins and a smaller fraction of TS/TE proteins (housekeeping adjusted *P*, synthesis vs. clearance:3.3e-8; folding vs synthesis: 1.9e-11; TS/TE: synthesis vs. clearance 0.017; MW). D. Expression uniformity across time and tissues per system. Protein synthesis was more uniformly expressed across time and tissues (adjusted *P* <5.6e-6; MW). E. Median expression levels in adult human tissues of each system. Protein synthesis was more highly expressed than protein folding, and protein folding was more highly expressed than protein clearance (MW test, adjusted *P* clearance vs. synthesis 1.2e-10; folding vs synthesis: 1.4e-6, folding vs. clearance: 1.4e-6). F. The impact on cell growth of each system. Protein synthesis had the largest impact on cell growth (adjusted *P* clearance vs. synthesis 1.7e-59; folding vs synthesis: 1.1e-26, folding vs. clearance: 0.005, one-sided KS test). G. The fraction of Mendelian disease genes per class. H. The age coefficient of mouse orthologs of genes per system. Protein synthesis increased more with age relative to other systems (KS test, adjusted *P*<0.0002).

## DISCUSSION

Protein clearance, a fundamental requisite in every living cell, is handled in eukaryotes by the UPS and ALP, which are highly conserved from yeast to humans. Parts of these clearance systems have been studied in depth, yet not much is known about their basal organization across tissues and cell types. Here, we harnessed recent and heterogeneous omics profiles of human tissues and cells to provide a better understanding of the properties and organization of the two clearance systems.

We first analyzed the expression ubiquity, stability, and essentiality of the two clearance systems in a variety of human tissues and cells and across time. These measures, along with conservation, were used previously to define housekeeping genes ^52,73–76^. In line with the fundamental roles of the two clearance systems, UPS and ALP components were more widely expressed across tissues and cell types in both adult and developing tissues, included more housekeeping proteins, and were more essential for growth across cell lines than other protein-coding genes. This supports the use of these omics-based measures to estimate the fundamental functionality of human molecular systems. When comparing the three branches of the proteostasis network, protein-folding behaved somewhat similarly to protein clearance, whereas protein synthesis was much more ubiquitous and essential. This resulted in a hierarchal organization where protein clearance is more fundamental than protein-coding genes and is superseded by protein-folding, with protein synthesis at the top of the hierarchy. This hierarchy suggests that although UPS and ALP contain fundamental elements, they possess a large degree of tissue plasticity.

UPS and ALP are relatively large systems, containing hundreds of genes belonging to distinct classes. Next, we zoomed into those classes. Most classes obeyed the general behavior observed above. Exceptions included the UPS E3 and the ALP lysosome classes, which were relatively more variable, and the UPS proteasome and the ALP ESCRT classes, which were significantly more fundamental. Interestingly, this was not related to the size of the class or to its role. For example, both E3 and the lysosome are highly variable, yet E3 genes compose 63% of the UPS (the largest class), while lysosome genes are only 14% of the ALP. Likewise, both the proteasome and the lysosome are endpoints of the clearance pathways, yet they show distinct behaviors.

UPS and ALP handle protein clearance in morphologically and functionally distinct tissues and cell types. To ask whether protein clearance systems are tailored to proteome demands of different cellular environments or act as ‘one size fits all’ systems, we analyzed their expression patterns in adult and developing human tissues and cell types. We found that most genes changed their expression across tissues. This was recently demonstrated for the 26S proteasome activity in *Arabidopsis thaliana* and mice revealing clear differences in proteasomal abundance and activity between tissues ^42,77,78^. We then asked whether the tissue-aware expression has functional relevance. Manual curation of genes upregulated in skeletal muscle showed that many of them were indeed associated with muscle function and disease. Furthermore, knockdown of muscle-upregulated genes not previously associated with muscle function resulted in reduced motility in *C. elegans* (Fig. 4C-D). Notably, tissues marked by upregulation of the protein clearance systems, such as brain and skeletal muscle tissues, were more susceptible to tissue-specific Mendelian diseases caused by aberrant protein clearance genes, such as familial Parkinson’s and Huntington’s diseases (Fig. 4B). Thus, the clearance systems are fundamental yet adaptable. We also found that UPS and ALP genes were more negatively correlated with age than other protein-coding genes in mice (Fig. 1E-F), in agreement with many protein misfolding diseases being late onset. We propose that tissue-specific differences in demand could expose the loss of essential functions and lead to tissue-specific phenotypes without other broad pathological impacts ^79^. For example, different mutations in UBA1, the main E1-ubiquitin-activating enzyme ^69^, cause tissue-specific diseases, including spinal muscular atrophy (SMAX2), adult-onset inflammatory disorder (VEXAS), and lung cancer (LCINS), that were linked to tissue-specific differences in E1-E2 dysregulation ^80^.

The expression of approximately 10% of the genes in each system was relatively uniform across tissues and cell types. We next asked if they serve as a core. Indeed, these genes were more highly expressed and essential relative to the variably expressed genes and included representatives from all the UPS and ALP classes. The existence of a core subset and the establishment of core-variable tissue-specific functional networks were previously observed in the chaperone system ^48,81,82^. We propose that core and variable organization is a basic feature of the proteostasis network, which allows it to carry basal cellular functions and yet respond to changing proteomic demands.

This study has several limitations. Firstly, the definition of the genetic components of the UPS and the ALP was not straightforward. UPS included not only ubiquitin but also other ubiquitin-like proteins such as SUMO and UFM1 and their associated E1 activating enzymes, E2 conjugating enzymes, and E3 ligases ^16^. Some genes function in multiple branches of the proteostasis network or in different classes of a specific branch; for example, PRKN is associated with ALP and UPS, whereas LRRK2 is associated with autophagophore initiation and elongation, autophagosome closure maturation and lysosome fusion and lysosomal catabolism. To define these systems and classes, we adopted the recent compilation of the proteostasis consortium, which associated some genes to more than one class ^35,36^, and validated associations using additional resources (see Methods). Secondly, much of our analyses used transcriptomic profiles, which were oblivious to post-transcriptional events. To confront this limitation, we incorporated proteomic profiles of 32 human tissues, which supported the main observations. We also included bulk profiles of 7 developing human tissues, single-cell RNA-sequencing profiles of 28 human tissues, essentiality analysis of human cell lines, and expression correlation with age in mice, to allow views across age and into cell types. Analyses of these datasets supported the fundamental role of each system and its organization into core and variable components. Lastly, our study focused on basal activities, yet systems change upon stress or other conditions. Nevertheless, correlation analysis of transcriptomes of 33 human cancers revealed behaviors that were similar to non-cancer samples. Taken together, we showed that proteostasis branches have a tissue-aware organization that responds to varied proteome demands and can thus contribute to tissue-selective phenotypes and diseases.

## MATERIALS AND METHODS

### List of human UPS, ALP, protein folding, and synthesis genes

We utilized the comprehensive enumeration of components of the human proteostasis system compiled recently by the proteostasis consortium ^35,36,49^. We followed the consortium’s classification of genes into branches, classes, and groups, whereby certain genes could belong to more than one branch, class, or group. UPS classes included E1 activating enzymes, E2 conjugating enzymes, E3 ubiquitin ligases, p97 and associated proteins, and proteasome and associated proteins. For simplicity, we united the additional UPS classes ‘ubiquitin and UBL like proteins’, ‘ubiquitin and UBL demodifiers’ and ‘ubiquitin and UBL binding’ under the collective class ‘ubiquitin and UBL’ (Table S1A). We removed two pseudogenes and a putative gene (FBL21, RP11-651P23.4, and SIK1, respectively) from the list of UPS genes. ALP classes included autophagy preinitiation signaling (including mTOR), gene expression, substrate selection, CDA, initiation and elongation, autophagosome closure maturation and lysosome fusion, lysosome reformation, and lysosome catabolism. For simplicity, the ALP class ‘lysosome reformation’ was merged with the class ‘autophagosome closure maturation and lysosome fusion’, as five of its nine genes overlapped with this class, except for the gene GBA which overlapped with the class lysosome catabolism. The merged class was collectively termed ‘maturation lysosome reformation and fusion’. Additionally, we defined the 27 genes of endosomal complexes required for transport (ESCRT), which were part of the lysosome closure and maturation class, as a distinct class (Table S1B). We removed three genes annotated as ‘specific function in autophagy unknown’ (PPARA, KIAA1324, MAP1LC3B2) from the list of ALP genes. The protein folding branch included the classes chaperone, folding enzymes, maturation and folding of specific substrates, and membrane protein folding (Table S1C). The protein synthesis branch included the classes ribosome, ribosome biogenesis factor, translation initiation, translation elongation, translation termination, and ribosome-associated QC (Table S1D). A set of protein-coding genes with unique ENSG IDs and HGNC names was extracted from ensemble-biomart ^83^ (Table S1E).

### Dataset of Mendelian diseases

Mendelian diseases and disease genes were obtained from the OMIM database ^84^, and limited to diseases with a known molecular basis (OMIM phenotype mapping key 3). Diseases arising from aberrations in UPS or ALP genes were associated with the tissues in which they manifested clinically based on OMIM records and literature curation ^85^.

### Analysis of bulk tissue transcriptomes

Bulk transcriptomic profiles of adult human tissues were obtained from the GTEx portal (GTEx v8). We analyzed samples from donors with a cause of death attributed to traumatic injury or suicide and a terminal phase estimated to last less than 10 minutes to limit the effect of medication and stress on the expression of proteostasis genes. Tissues with less than five samples were removed from the analysis. As in ^86^, brain sub-regions were grouped into six regions: Cortex (including anterior cingulate cortex (BA24), hippocampus, cortex, frontal cortex); cerebellum (including cerebellum, cerebellar hemisphere); basal ganglia (including caudate, nucleus accumbens, putamen); spinal cord; hypothalamus; and amygdala. The resulting dataset included 665 samples of 34 tissues. Genes were considered as expressed in a tissue if their median expression in samples of that tissue was ≥ 5 transcripts per million (TPM). In analyses of expression in adult human tissues, only genes that were considered as expressed in at least one tissue were included. In total, we included 1127 UPS, 812 ALP, 294 protein folding, 324 protein synthesis genes, and 16,771 other protein-coding genes. To assess the expression ubiquity of genes across tissues, we associated each gene with the number of tissues expressing it.

### Differential gene expression analysis

Differential gene expression per tissue relative to other tissues was calculated as in ^48^. Specifically, raw reads were normalized to obtain the same library size per sample by using the edgeR TMM method ^87^ and transformed using voom ^88^. Genes with ≤10 raw counts per sample across all samples were removed before normalization. Differential gene expression per tissue was computed by applying the Limma linear model to the transformed profiles of that tissue and to a background set containing the transformed profiles of all other tissues. The differential expression of a gene in a tissue was set to its log_2_ fold-change value. *P*-values were adjusted for multiple hypothesis testing via Benjamini–Hochberg correction. We considered genes with an absolute log_2_ fold-change≥1 and adjusted *P*-value≤0.05 as differentially expressed. We defined core genes as genes that were expressed in all tissues above 5TPM with an absolute log_2_ fold-change<1. All other genes were defined as variably expressed.

### Analysis of single-cell tissue transcriptomes

Single-cell RNA-sequencing data from 28 adult human tissues were obtained from Tabula Sapiens ^51^. Analysis was performed using Seurat package v4.0.5 ^89^. Per tissue, gene expression levels were normalized cell-wise using the NormalizeData function. We considered only genes with normalized counts ≥0.05 in at least 10% of all cells. To limit noise, a gene was considered expressed in a cell type if its expression level was ≥ 0.15 normalized counts. In total, data were available for 1078 UPS, 820 ALP, 290 protein folding, 323 protein synthesis genes, and 14,148 other protein-coding genes. To assess the expression ubiquity of genes across cell types per tissue, we associated each gene with the number of cell types expressing it.

### Tissue-specificity of proteins

Proteomic profiles of 32 human tissues were obtained from ^52^. As described therein, proteins were assigned tissue specificity scores and classified as tissue-specific (TS), tissue-enriched (TE), housekeeping, or other. Specifically, TS proteins had a tissue specificity score ≥ 4 in one tissue and no score > 2.5 in any other tissue. TE proteins were (i) expressed in at least three tissues, (ii) had a tissue specificity score ≥ 2.5 in at least one tissue, and (iii) were not tissue-specific. Housekeeping proteins had tissue specificity scores <2 in every tissue. All other proteins were annotated as ‘other’. For simplicity, we combined TS and TE proteins into one group, denoted TS/TE. In total, data were available for 809 UPS, 706 ALP, 268 folding, 315 synthesis proteins, and 10,271 other proteins.

### Organ and timepoint specificity of genes

Data of organ and timepoint specificity of genes was extracted from ^55^, which conducted bulk transcriptomic analysis of seven organs at 23 time points, ranging from early organogenesis (14 time points in weeks 4-20 post-conception) to adulthood. Organ specificity reflected whether a gene was expressed highly in a single organ. Timepoint specificity per organ reflected whether a gene was expressed highly at a single time point. Specificity values ranged from 0 (non-specific) to 1 (most specific). For clarity, we transformed specificity to uniformity, which we defined as 1 minus specificity. Data were available for 1132 UPS, 808 ALP, 291 folding, 326 synthesis, and 14,648 other protein-coding genes.

### Analysis of bulk cancer transcriptomes

Bulk transcriptomic profiles of human cancers were obtained from the cancer genome atlas (TCGA). To test for expression changes we calculated, per cancer, the preferential expression of the gene in that cancer relative to all other cancers, according to ^90^ (Equation 1). Genes with a preferential expression ≥2 were defined as preferentially expressed in that cancer. Data were available for 1068 UPS, 776 ALP, 281 folding, and 303 synthesis genes.

Equation 1:

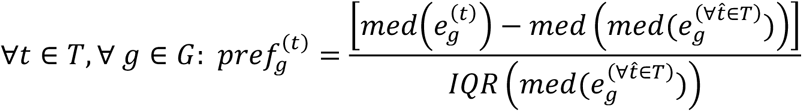

*T* denotes the set of tissues; *G* denotes the set of genes; *pref* denotes preferential expression; *e* denotes normalized count; *IQR* denotes inter-quartile range. For any tissue *t*, 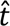 is any other tissue in *T*.

### Gene essentiality scores

Data on gene essentiality scores was obtained from DepMap ^56^ via the DepMap portal ^91^ and included CRISPR scores for 17,453 genes in 1078 cell lines. A negative (or positive) CRISPR score of a gene in a cell line implied a reduced (or increased) growth rate of a cell line containing the CRISPR-inactivated gene relative to control. We set the essentiality score of a gene to its minimal CRISPR score across cell lines. Data were available for 1062 UPS, 673 ALP, 283 folding, 280 synthesis genes, and 13,717 other protein-coding genes.

### Correlation with age

Data of age coefficient per tissue and gene were obtained from *Tabula senis* ^92^. We associated each human gene with its homologous mouse gene using data on orthologous genes from Mouse Genome Informatics (MGI) website ^93,94^. Data were available for 999 UPS, 736 ALP, 268 folding, 274 synthesis genes, and 12,670 other protein-coding genes. To compare gene subsets across tissues, each gene was associated with its median age coefficient across tissues.

### Curation of genes with muscle-associated functionality

We compiled lists of the top 80 muscle-upregulated UPS and ALP genes (Table S3). We annotated the muscle-associated functionality of these genes by manual curation of their OMIM entries and by literature search. For each gene, we noted the Mendelian diseases it was linked to and its association with muscle function (Table S3).

### Nematode orthologs

We used the OrthoList 2 tool for comparative genomic analysis between *C. elegans* and humans ^95^ to identify *C. elegans* orthologs of muscle-upregulated human genes with no previously known muscle phenotypes.

### Nematodes cultivation and feeding RNAi

N2 (wild-type) nematodes were grown on NGM plates seeded with the *Escherichia coli* OP50-1 strain and maintained at 20°C. Age-synchronized embryos were obtained by placing ∼40 adults for 1 hour on RNAi plates seeded with *E. coli* strain HT115(DE3) transformed with empty vector control (pL4440) or RNAi vectors for *M60.7* (human ASB15), *B0545.1/tpa-1* (PRKCQ), *T27A3.2/usp-5* (USP13), *F10D7.5* (NEURL1), and *T01C8.1/aak-2* (PRKAA2) obtained from the Ahringer RNAi library ^96^.

### Motility assay

Motility rates were determined by counting the number of changes in bending direction at mid-body (thrashes) per minute of 20 animals per treatment ^97^. Animals were scored on the first day of adulthood after the onset of egg-laying.

### Statistical analysis

To test the null hypothesis that two distributions were drawn from the same continuous distribution, we used the Kolmogorov-Smirnov test (relevant to the distribution of expressed genes across one, two, or more tissues and to the distribution of gene’s essentiality scores). To test the null hypothesis that two sets had similar values, we used the Mann-Whitney U test (relevant to gene expression levels per tissue and to the number of cell types expressing the gene per tissue). To test the null hypothesis that the overlap between two sets was similar to the expected, we used the Fisher exact test (relevant to overlap between tissue disease genes and tissue upregulated genes). Per analysis, *P*-values were adjusted for multiple hypothesis testing using the Benjamin-Hochberg method. Correlation between fractions of up- or down-regulated genes was computed using Pearson correlation. Finally, we used one-way ANOVA followed by Dunnett’s post-hoc test to compare motility rates between control and RNAi-treated animals.

## Acknowledgments

This study was funded by the Israel Science Foundation [317/19 and 401/22 to E.Y.-L. and 278/18 and 420/23 to A.B.-Z].

## Author contributions

Conceptualization, A.B.-Z., and E.Y.-L.; Formal analysis, E.V., L.R., J.J.; Investigation, E.V., L.R., and B.E.; Writing—original draft, E.V., A.B.-Z., and E.Y.-L.; Supervision, A.B.-Z. and E.Y.-L; Funding acquisition, A.B.-Z. and E.Y.-L.

## Competing interests

The authors declare no competing interests.

## Notes

### Competing Interest Statement

The authors have declared no competing interest.

