## Supplementary material for "The landscape of cellular clearance systems across human tissues and cell types is shaped by tissue-specific proteome needs": Supp Figures 1-4

**Vinogradov, et al**

Supplementary Figures 1-4

### Supplementary Figures

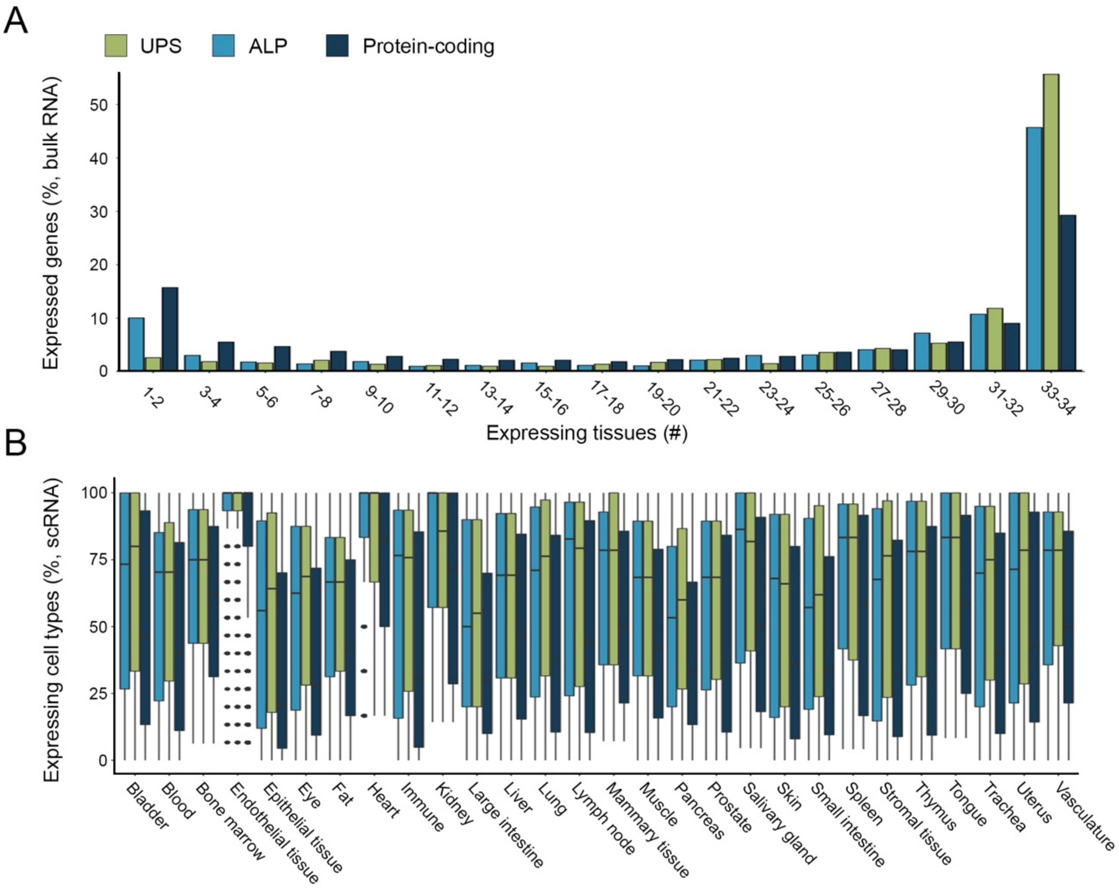


**Figure S1. Expression patterns of UPS, ALP and other protein-coding genes across adult human tissues.**

A. The number of adult human tissues that express UPS, ALP, and other protein-coding genes. Analysis was based on bulk RNA transcriptomes of 34 tissues. Data included 1127 UPS, 812 ALP, and 16,771 other protein-coding genes that were expressed in at least one tissue. UPS and ALP genes were expressed in more tissues than other protein-coding genes (KS test, adjusted *P* UPS vs. protein-coding: 0.0086; ALP vs protein-coding 3.1e-4).

B. The expression levels of UPS, ALP, and other protein-coding genes per tissue. UPS and ALP had higher expression levels than other protein-coding genes. Data included 1127 UPS, 812 ALP, and 16,771 other protein-coding genes that were expressed in at least one tissue; genes were included in a given tissue only if they were expressed in that tissue.

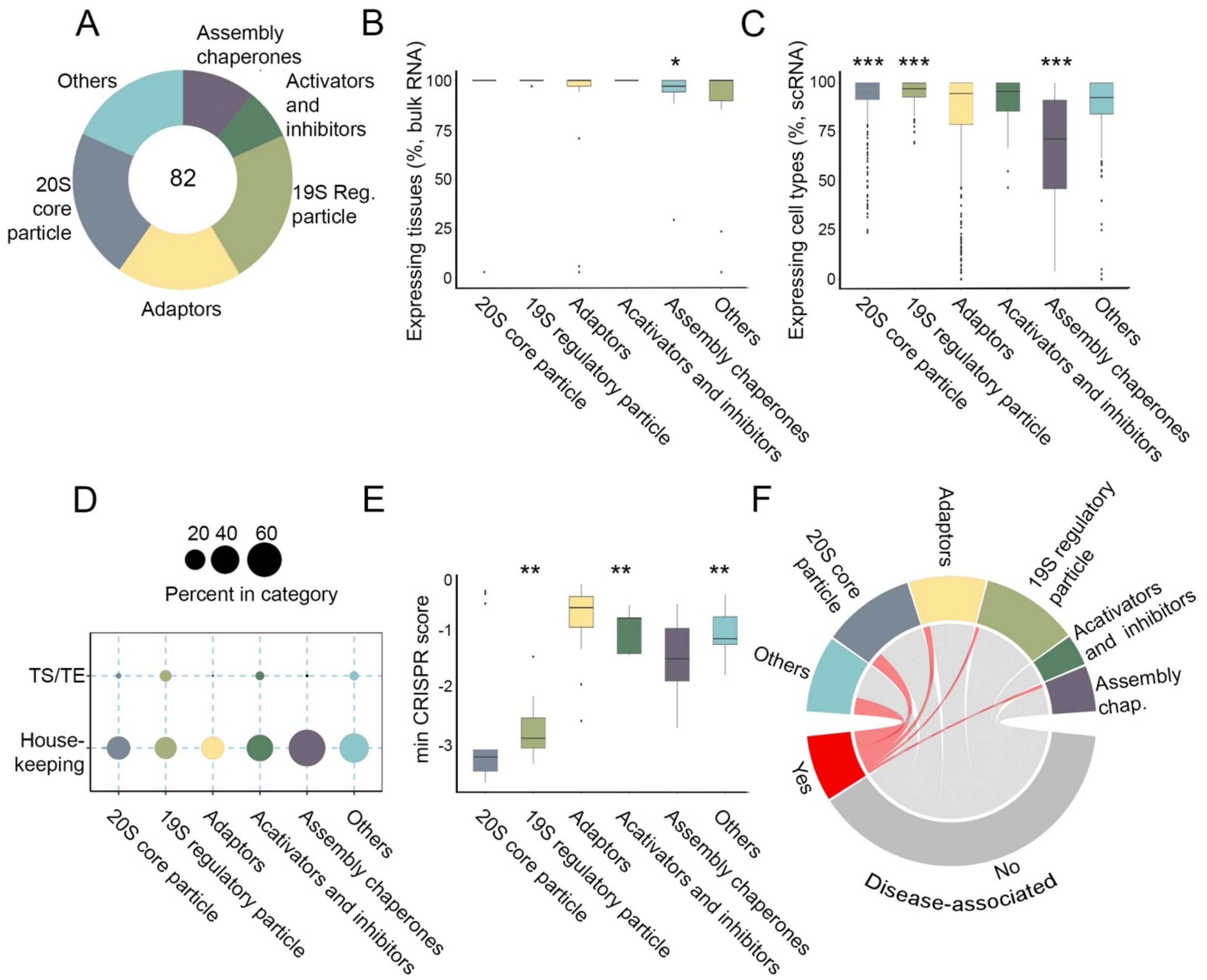


**Figure S2: Expression patterns and phenotypic impact of proteasome groups.**

A. The relative number of genes per proteasome groups: 20S core particle, 18; 19S regulatory particle, 19; adaptors, 15; activators and inhibitors, 6; others, 15.

B. The percent of adult human tissues that express proteasome genes per group. 20S core and 19S regulatory particles were expressed in all tissues (adjusted *P*=0.06 and 0.07, respectively; MW).

C. The percent of cell types per tissue that express proteasome genes per subclass. 20S core and 19S regulatory particles were the most ubiquitous (adjusted *P*=1e-08 and 2.5e-19, respectively; MW).

D. The fraction of housekeeping and TS/TE proteins per proteasome subclass. 20S core and 19S regulatory particles had similar factions of housekeeping genes, and the 19S regulatory subunits had a larger fraction of TS/TE genes.

E. The impact on growth of proteasome groups. 20S core and 19S regulatory particles had the highest impact on cell growth (adjusted *P*=5e-4, MW).

F. The fraction of genes associated with Mendelian diseases per proteasome subclass. Activators and inhibitors genes were not associated with Mendelian diseases.

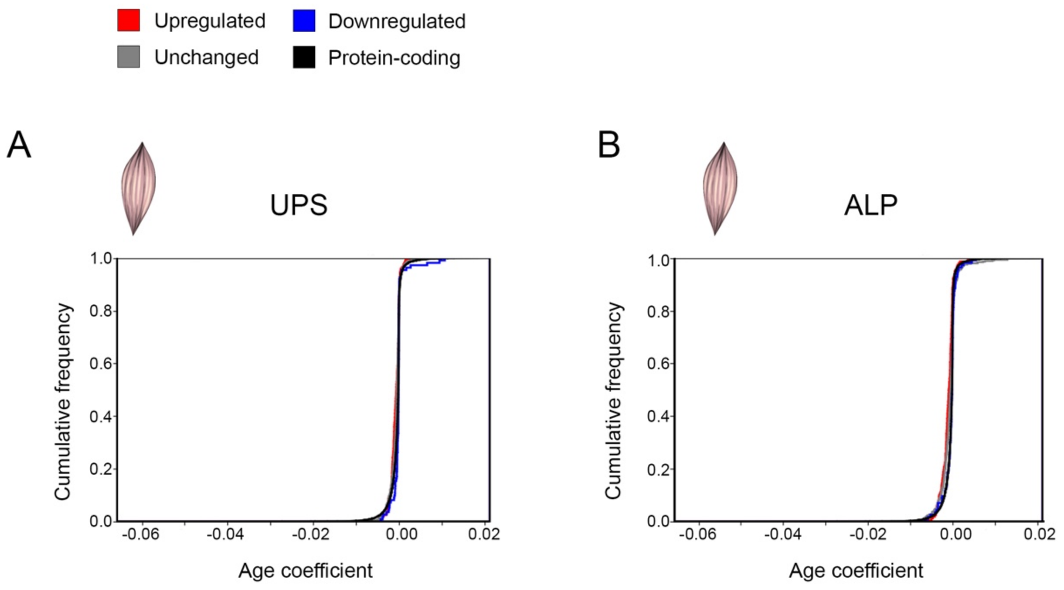


**Figure S3. The age coefficient in limb muscle of mouse orthologs of UPS and ALP genes that were upregulated, unchanged, or downregulated in human skeletal muscle tissue.** The numbers of human UPS and ALP genes with age coefficient data were 590 and 456 unchanged, 279 and 159 upregulated, and downregulated 110 and 113, respectively. In UPS, upregulated genes had lower age coefficients than unchanged genes (KS test, adjusted *P*=0.004), and both had lower age coefficients relative to downregulated genes (KS test, adjusted *P* <9e-10 and 6.1e-11, respectively). In ALP, upregulated genes and unchanged genes had lower age coefficients than downregulated genes (KS test, adjusted *P*<7.4e-8 and 0.0001, respectively).

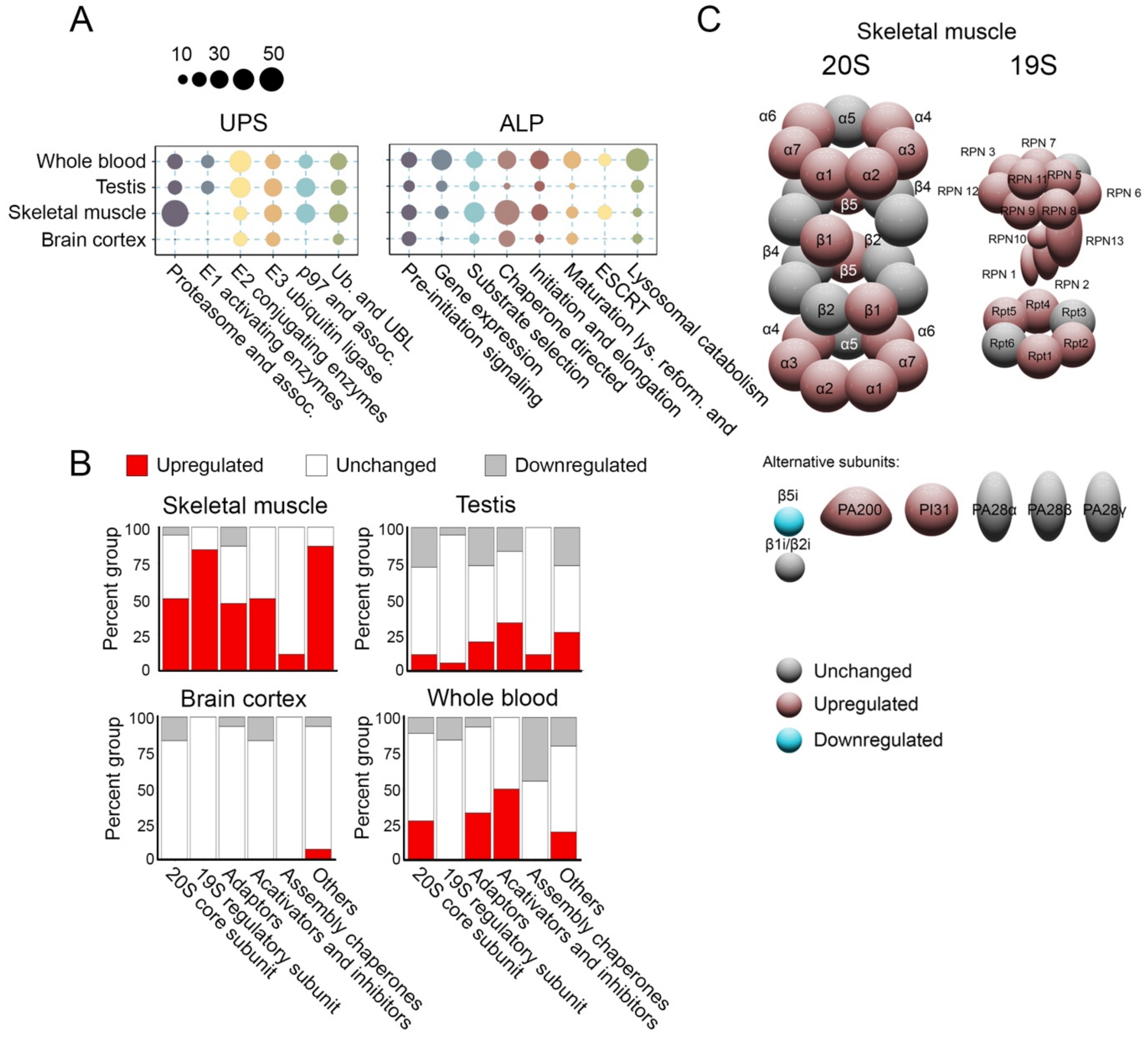


**Figure S4. UPS and ALP classes and groups are variably expressed in skeletal muscle, brain cortex, testis, and whole blood.**

A. The fractions of upregulated genes per class in adult human tissues (expression fold-change ≤2). Proteasome and CDA had larger fractions of upregulated genes in skeletal muscle tissue.

B. The percent of genes that were upregulated, downregulated, or unchanged in proteasome groups per tissue. The largest fractions of upregulated genes were in skeletal muscle tissue, and almost all genes were unchanged in brain cortex tissue.

C. A schematic representation of upregulated (red), downregulated (teal), and unchanged (gray) genes in the proteasome 20S and 19S particles in skeletal muscle tissue. Most of the 19S genes were upregulated.
